# Decoding host responses: How a freshwater invertebrate defends against a parasitic bacterium

**DOI:** 10.1101/2025.03.28.645445

**Authors:** Gattis Sabrina, Paraskevopoulou Sofia, Marcus Jade Rachel, Ben-Ami Frida

**Affiliations:** School of Zoology, George S. Wise Faculty of Life Sciences, Tel Aviv University, Tel Aviv, 6997801, Israel; Division of Molecular Biosciences, Department of Biology, University of Lund, Lund, Sweden

**Keywords:** Host-parasite interactions, *Daphnia magna*, *Pasteuria ramosa*, immune response, RNA-seq

## Abstract

Host-parasite interactions drive coevolutionary dynamics, often leading to reciprocal adaptations between hosts and parasites, a process known as the Red Queen arms race. While the molecular mechanisms underlying vertebrate and insect immune responses have been studied extensively, those of aquatic invertebrates remain unexplored, despite their critical role in ecosystem stability and aquaculture. Here, we use the *Daphnia magna–Pasteuria ramosa* system to investigate the host’s immune response and the molecular mechanisms underlying host-parasite interactions. We inoculated 800 *D. magna* hosts with 20,000 mature spores of *P. ramosa* and tracked the progression of infection by measuring the proportion of infected individuals and the developmental stages of the parasite at multiple time points post-inoculation (hereafter p.i.). RNA sequencing was performed at key infection phases, early (Cauliflower stage), mid (Cauliflower and Grape stages), and terminal (Cauliflower, Grape, and Mature spore stages), to capture gene expression changes linked to infection dynamics. Our transcriptomic analyses revealed key immune genes involved in host defense, including genes involved in Toll signaling pathways, thereby revealing significant changes in pathways related to immune function, host metabolism, and resource allocation. Our findings suggest that iron sequestration may serve as a host defense strategy to restrict parasite growth, representing a form of nutritional immunity. Furthermore, pathways associated with infection-induced phenotypic traits, such as somatic growth, red coloration, and castration, were significantly upregulated, underscoring the impact of the infection on host physiology. Taken together, these findings provide new insights into the interplay between hosts and parasites at a molecular level in an ecologically relevant system, advancing our understanding of infection strategies in aquatic invertebrates.

## Introduction

Host-parasite interactions impose strong selective pressures, often leading to the reciprocal adaptation between both antagonists, a process known as co-evolution (reviewed in Langerhans, 2008). One of the most critical aspects of this evolutionary arms race is the host’s ability to detect and counteract infections, primarily through its immune system (Schmid-Hempel, 2009). While vertebrates possess a sophisticated adaptive immune system characterized by immunological memory (Litman et al., 2010), invertebrates rely on their innate immune system to combat parasitic infections (reviewed in Coates et al., 2022). This innate immune system includes a diverse set of cellular and molecular defenses, such as receptors that recognize pathogen-associated molecular patterns (PAMPs), regulators that modulate signaling pathways, e.g., the “Toll pathway”, and effectors that are directly engaged in the inhibition of pathogen proliferation or survival (Coates et al., 2022; Schlenke & Begun, 2004; Schmid-Hempel, 2009).

The progression of a parasitic infection is influenced largely by how the parasite develops within the host, as different stages impose distinct physiological and immunological challenges. When an immune response is triggered, specific genes involved in pathogen defense, such as those encoding for antimicrobial peptides, lysozymes, and lipoproteins, are upregulated to counteract the infection (reviewed in Coates et al., 2022). As the infection progresses, parasites employ different strategies to evade host immune defenses. In response, hosts adjust their defensive mechanisms to counteract the effects of the parasites, e.g., via nutrient regulation. Several hemoglobin-binding proteins in vertebrates capture the free hemoglobin to prevent oxidative damage and deprive pathogens of accessible iron (Ong et al., 2006). In invertebrates, homologues of iron-sequestering proteins, including transferrin and ferritin, have been associated with innate immunity (Ong et al., 2006). In insects, iron-binding proteins of the transferrin family frequently play a prominent role in iron sequestration during infection (Hrdina & Iatsenko, 2022). Another defense mechanism involves resource allocation to immune response, e.g., hosts may divert energy and resources from growth and reproduction to bolster their immune defenses (Raw, 2012). Finally, during later phases of infection, the host immune response may shift from pathogen elimination towards repair and metabolic adjustments. *Drosophila melanogaster*, for instance, suppresses excessive immune activation to minimize self-damage, while maintaining physiological function (Ayres & Schneider, 2012).

While most immunological research has focused on vertebrates and insects, non-insect arthropods, such as *Daphnia*, can provide key insights into the diversity of immune defensive mechanisms in organisms that lack adaptive immunity. In freshwater ecosystems, *Daphnia* plays a critical ecological role as a keystone grazer, controlling phytoplankton populations, while serving as prey for higher trophic levels (Ebert, 2022). Natural *Daphnia* populations are frequently exposed to a diverse range of pathogens, including bacteria, fungi and microsporidians, which can have significant consequences for host fitness (Ebert, 2005; Goren & Ben-Ami, 2013). Among these, the bacterial parasite *Pasteuria ramosa* has been particularly well studied due to its severe effects on host survival and reproduction. Upon infection, *P. ramosa* invades the hemolymph, proliferates within the host, and eventually induces host castration, leading to population declines that may disrupt entire food web structures (Ebert et al., 2004). Therefore, understanding host-parasite interactions in *Daphnia* is crucial for predicting how disease outbreaks affect the resilience of freshwater ecosystems.

Despite the ecological importance of these animals, the molecular mechanisms underlying their responses to infection at different phases remain poorly understood. Previous studies have identified candidate immune genes involved in pathogen defense (e.g., Phenoloxidase; Labbé & Little, 2009) and have shown that *Daphnia magna* exhibits a rapid and dynamic transcriptional response upon *P. ramosa* exposure (McTaggart et al., 2015). Further transcriptomic analyses in other *Daphnia* species have reinforced the idea that host immune responses may shift throughout infection, depending on parasite burden and developmental stage. For example, in *Daphnia galeata* a downregulation of genes related to immune function and metabolism was detected 48 hours post-infection with microsporidia, suggesting a trade-off between immune defense and metabolic homeostasis (Lu et al., 2018). Similarly, *D. dentifera* infected with the yeast *Metschnikowia bicuspidata* exhibited an increase in the number of differential expressed genes as infection progressed, and significantly enriched Gene Ontology (GO) terms were related to cuticle development and defense responses (Terrill Sondag et al., 2023).

These findings suggest that *Daphnia’s* immune responses are highly dynamic, varying not only across different host genotypes and parasite species, but also throughout the course of infection. Here, we employed RNA-sequencing (RNA-seq) to investigate how gene expression in *D. magna* varies at distinct time points representing key phases of *P. ramosa* infection. By analyzing gene expression across infection process where different parasite developmental stages are present, we aim to identify host responses associated with parasite establishment, persistence and proliferation, thereby providing valuable insights into how *Daphnia* modulates its immune response over the progression of the infection.

## Material and methods

### Experimental design

*Daphnia* individuals (HO2 clone, Hungary) were acclimated in standardized lab conditions (20±1°C, 16:8 L:D) for two generations. *Daphnia* for the experimental generation were isolated from the 3^rd^ brood of the acclimation generation, and fed daily according to an age-dependent feeding protocol (Izhar & Ben-Ami, 2015) to provide *ad libitum* resources. Five days post-birth 800 juveniles were assigned to two treatments: animals inoculated with the parasite (Treatment I) and animals without the parasite, i.e., the control treatment (Treatment C). The larger initial number of animals accounted for random mortality and ensured an ample sample size for random sampling until the final collection point. During inoculation, each individual was placed in jars containing 20 mL of Artificial *Daphnia* Medium (ADaM; Ebert et al., 1998; Klüttgen et al., 1994) and exposed to 20,000 mature spores of *P. ramosa* (clone C19), if belonging to the parasite group. For the control group, the same amount of a placebo solution composed of crushed uninfected *D. magna* was administered. *Daphnia* were monitored daily for offspring production and offspring were removed, throughout the inoculation period, to minimize dilution effects. Seven days p.i., all *Daphnia* were transferred to jars filled with 80 mL of fresh ADaM, and thereafter until the end of the experiment, medium was replaced three times a week.

### Infection rate and spore stage production

On days 8, 11, 13, 15, 25 and 35 p.i., 20 individuals from each treatment group were randomly sampled and stored at -20°C in 100 μL ADaM for assessing infection and determining the presence of the three developmental stages of the parasite: “Cauliflower” spore stage (the first visible parasite developmental stage), “Grape” spore stage (the second visible parasite developmental stage), and “Mature” spore stage (the last parasite developmental stage, which is released into the environment upon host death). To assess infection rate, and determine spore concentration and developmental stage, *Daphnia* were crushed, and parasite stages were observed and counted using a Thoma cell counting chamber under a phase contrast microscope (Leica DM 2500). Infection rate was analyzed with binary logistic regression (R *glm* function, family=binomial), with “time post-inoculation” as the main effect. Post-hoc comparisons were performed with the “emmeans” R-package (Lenth, 2016), and data were visualized with the “ggplot2” R-package (Wickham, 2016). At each time point p.i., we recorded the number of infected hosts in which Cauliflower, Grape or Mature spores were detected. If an individual harbored multiple parasite developmental stages, it was counted in each corresponding category.

### RNA isolation and library preparation

On days 8, 11, 13, 15, 25 and 35 p.i., 20 individuals from each treatment group were sampled, homogenized with a sterile plastic pestle, and stored at -80°C in 500 μL Trizol. Based on the results of the phenotypic assay (see *Results* section), we selected three representative time points, 11, 15 and 35 days p.i., for RNA-sequencing. For each selected time point, four *Daphnia* replicates were used. Prior to RNA extraction, 10 μl of each replicate was examined under a phase-contrast microscope to confirm its infection status and determine the relevant developmental stages. Thus, our experimental design consisted of the following treatments, where inoculated hosts at each infection phase were compared to their respective uninfected controls at the same time point: (1) Early infection (I-Early *vs.* C-Early) – infected hosts 11 days p.i., harboring only Cauliflower spores, were compared to uninfected controls. (2) Mid infection (I-Mid *vs.* C-Mid) – infected hosts 15 days p.i., harboring both Cauliflower and Grape spores, were compared to uninfected controls. (3) Terminal infection (I-Terminal *vs.* C-Terminal), – infected hosts 35 days p.i., harboring all the three infection stages (Mature, Cauliflower and Grape spores), were compared to uninfected controls.

Subsequently, RNA was extracted with a combination of Trizol and column precipitation by using the RNeasy® Mini Kit (QIAGEN GmbH, Hilden, Germany) and applying a double elution step of 20 µl. RNA quantity and quality were confirmed with Qubit (Thermo Fisher Scientific) and 2200 TapeStation (Agilent, Santa Clara, CA), respectively. For library preparation, mRNA was enriched from 300 µl of total RNA with a Poly(A) enrichment protocol (NEBNext® Poly(A) mRNA Magnetic Isolation Module, New England Biolabs) and strand-specific libraries produced using NEBNext® Ultra™ II Directional RNA Library Prep Kit (New England Biolabs). Single-end (SE) sequencing of 100 bp reads was performed on a NovaSeq6000 Illumina platform (S1 Flowcell) by the Crown Genomics institute of the Nancy and Stephen Grand Israel National Center for Personalized Medicine (Weizmann Institute of Science, Rehovot, Israel).

### Differential gene expression analysis

Raw reads were trimmed with Trimmomatic 0.39, and sequence quality was determined before and after trimming with Fastqc (Andrews, 2010). Trimmed reads were mapped to the *D. magna* genome (daphmag2.4; GCA_001632505.1) with STAR (Dobin et al., 2013) and gene-level quantification estimates were produced by RSEM (Li & Dewey, 2011) (Table S1). Data were imported into R/Bioconductor with the “Tximport” R-package (Soneson et al., 2015) and gene expression was estimated using the “DESeq2” R-package, which performs a size factor normalization to account for differences in sequencing depth among libraries (Love et al., 2014). Genes with zero counts across all samples and those with low counts (< 10) in less than 25% of the samples were pre-filtered. To reveal insights into parasitic infection development, differential gene expression analyses were performed separately in three pairwise comparisons: 1) I-Early vs. C-Early, 2) I-Mid vs. C-Mid, and 3) I-Terminal vs. C-Terminal. Genes were considered differentially expressed at the level of 1.5-fold change (LFC threshold set to 0.58) and a p-adjusted value <0.05. All differentially expressed genes (DEGs) were further annotated against the NCBI non-redundant (*nr*) database (downloaded on the 15 March 2024) using the *blastx* algorithm with an e-value cutoff of 1e−10, and further annotated with GO Terms with OmicsBox version 3.0.30 (Conesa et al., 2005; Götz et al., 2008) (Table S2). Enrichment analyses were conducted in OmicsBox using Fisher’s exact test to assess the overrepresentation of specific biological functions in target gene sets. To account for multiple testing, the p-values were adjusted using the Benjamini-Hochberg method (Al-Shahrour et al., 2004). Overrepresentation was inferred at a False Discovery Rate (FDR) <0.05.

### Differential Pathway-Level expression analysis

To assess gene functions at the pathway level, all annotated genes were assigned KEGG Orthology (KO) terms using the online Kyoto Encyclopedia of Genes and Genomes (KEGG) automatic server (Kanehisa & Goto, 2000). To determine significant changes in pathway-level gene expression, we performed a pathway expression analysis using the “GAGE” v.2.46.1 R-package (Luo et al., 2009), which identifies differentially regulated pathways by evaluating expression changes across entire gene sets, rather than single genes. A stringent false discovery rate (FDR) cutoff of <0.01 was applied to minimize false positives. Additionally, redundancy was reduced by prioritizing pathways with the most distinct and functionally relevant gene contributions, ensuring a clearer biological interpretation, using the *esset.grp* function in the “GAGE” R-package. The R-package “Pathview” v. 3.5.1 (Luo & Brouwer, 2013) was used for visualization.

## Results

### Infection rate and parasite spore development

Infection rate was significantly affected by “time post-inoculation” (LR=97.34, p<0.001; Figure 1A), with the number of infected individuals increasing over time (post-hoc: 8 vs. 13: p=0.037; 8 vs. 15: p<0.016; 8 vs. 25: p<0.003; 8 vs. 35: p<0.002; 11 *vs*. 13: p<0.045; 11 *vs*. 15: p=0.01; 11 *vs.* 25: p=0.001; 11 *vs.* 35: p=0.006). Over time, we observed the accumulation of all three parasite stages. Specifically, the “Cauliflower” stage first appeared at 11 days p.i. in 15% of the animals and persisted until day 35 in 44% of the animals. The “Grape” stage was observed in 5% of the animals already by day 11 p.i. and reached 100% by day 35. From day 25 onwards, “Mature” spores were observed in 60% of the young animals, increasing to 84% at 35 days p.i. (Figure 1B).

**Figure 1.**
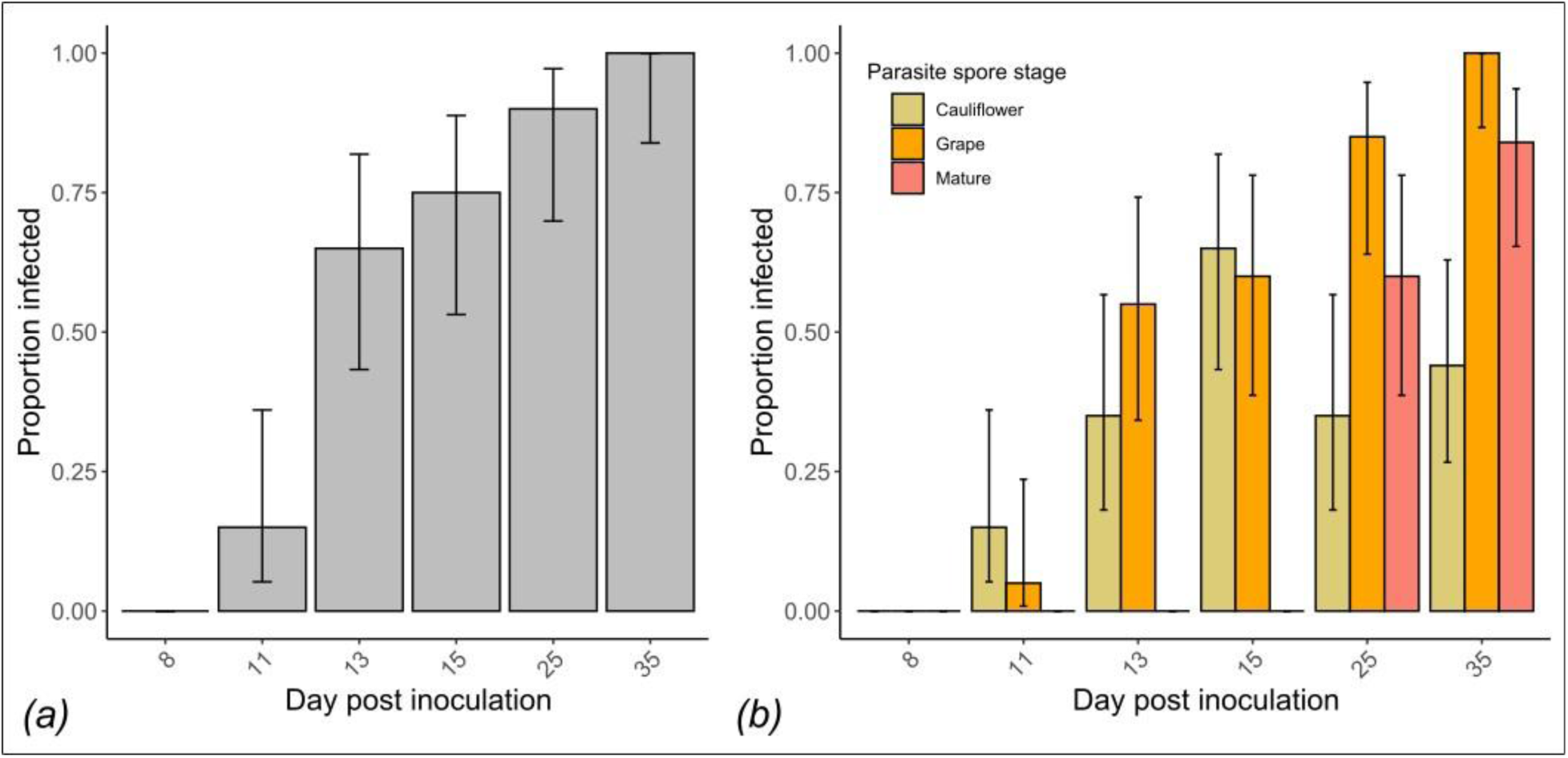
*(a)* Proportion of infected *D. magna* hosts at different time points post-inoculation. *(b)* Proportion of parasite developmental stages (i.e., “Cauliflower”, “Grape” and “Mature” spores) in *D. magna* at different time points post-inoculation. In both panels, error bars represent the 95% Wilson Confidence Interval.

### Across infection phases differential gene expression analysis

To gain insights into gene-specific expression associated with infection phase in *D. magna* hosts, we conducted pairwise comparisons between uninfected and infected animals during the Early, Mid, and Terminal infection phases. Comparing uninfected (C) to infected (I) hosts during the Early infection phase, we identified 567 DEGs (353 upregulated & 214 downregulated in infected hosts; Figure 2; Table S3). During the Mid infection phase, we detected 105 DEGs (47 upregulated & 58 downregulated in infected hosts; Figure 2; Table S4). Finally, during the Terminal infection phase, we detected 518 DEGs (313 upregulated & 205 downregulated in infected hosts; Figure 2; Table S5).

**Figure 2.**
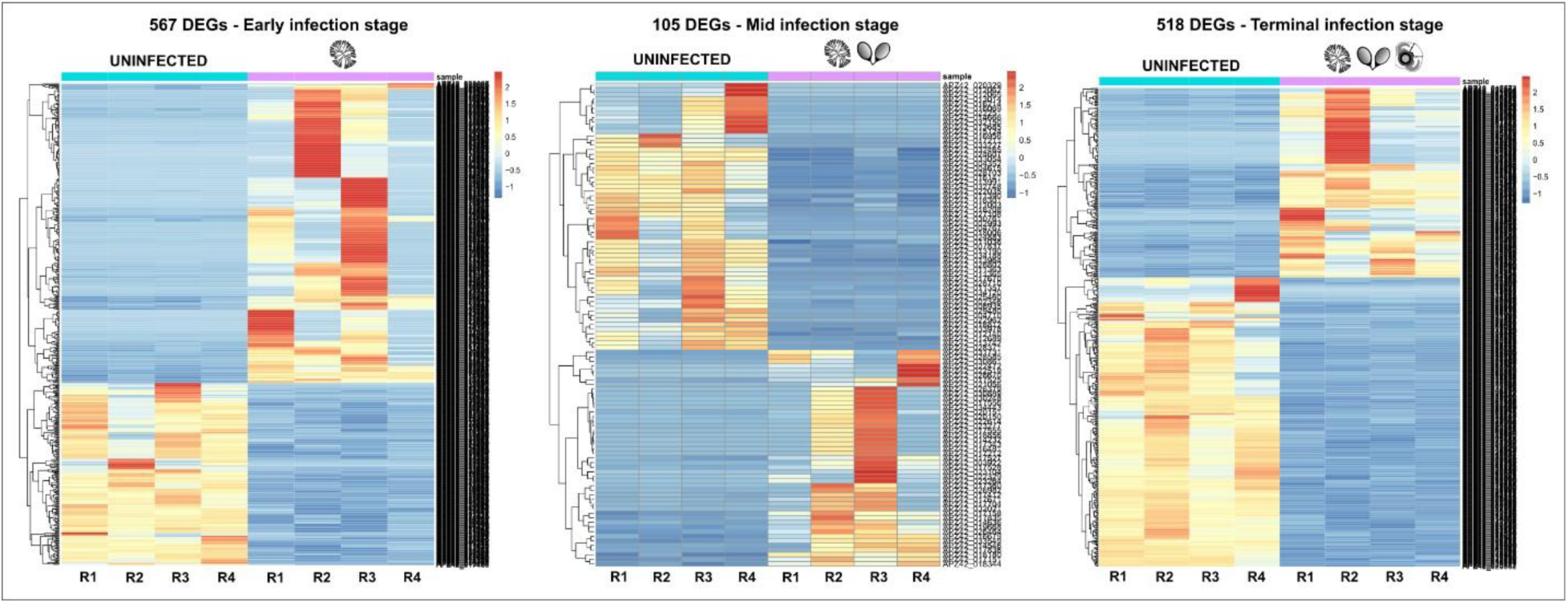
Heatmaps showing significant differentially expressed genes (DEGs) based on normalized counts (normalization performed DESeq2) for all pairwise comparison across infection phases. Upregulated genes are represented by the red spectrum, while downregulated genes are represented by the blue spectrum. Color bars represent the z-score.

### Across infection phases functional enrichment analyses

Thirty-three (33) GO terms were enriched during the Early infection phase, while 42 during the Terminal infection phase (Figure 3; Tables S6, S7). In the Mid infection phase, we did not yield any enriched functions. Twenty-nine (29) functions were enriched in both Early and Terminal infection phases, primarily involving gene regulation, transcriptional activity and structural components, thus indicating a continuous host response throughout infection. Six functions were uniquely enriched in the Early infection phase, among which are the “Chitin binding; GO:0008061” and “iron ion binding; GO:0005506” GO terms (Figure 3, Table S6). Genes associated with these GO terms included those encoding for cuticular proteins, endocuticle glycoproteins, N-acetylglucosaminyltransferases and chitin-based proteins for cuticle development. Fifteen (15) functions were uniquely enriched in the Terminal infection phase (Figure 3, Table S6). Among them the “Heme binding; GO:0020037” GO-term was enriched and genes encoding for Cytochrome P450 and Hemoglobin were upregulated. None of the enriched GO terms belonged to the “Immune response; GO:0009605” or the “Immune system process; GO:0002376” GO terms.

**Figure 3.**
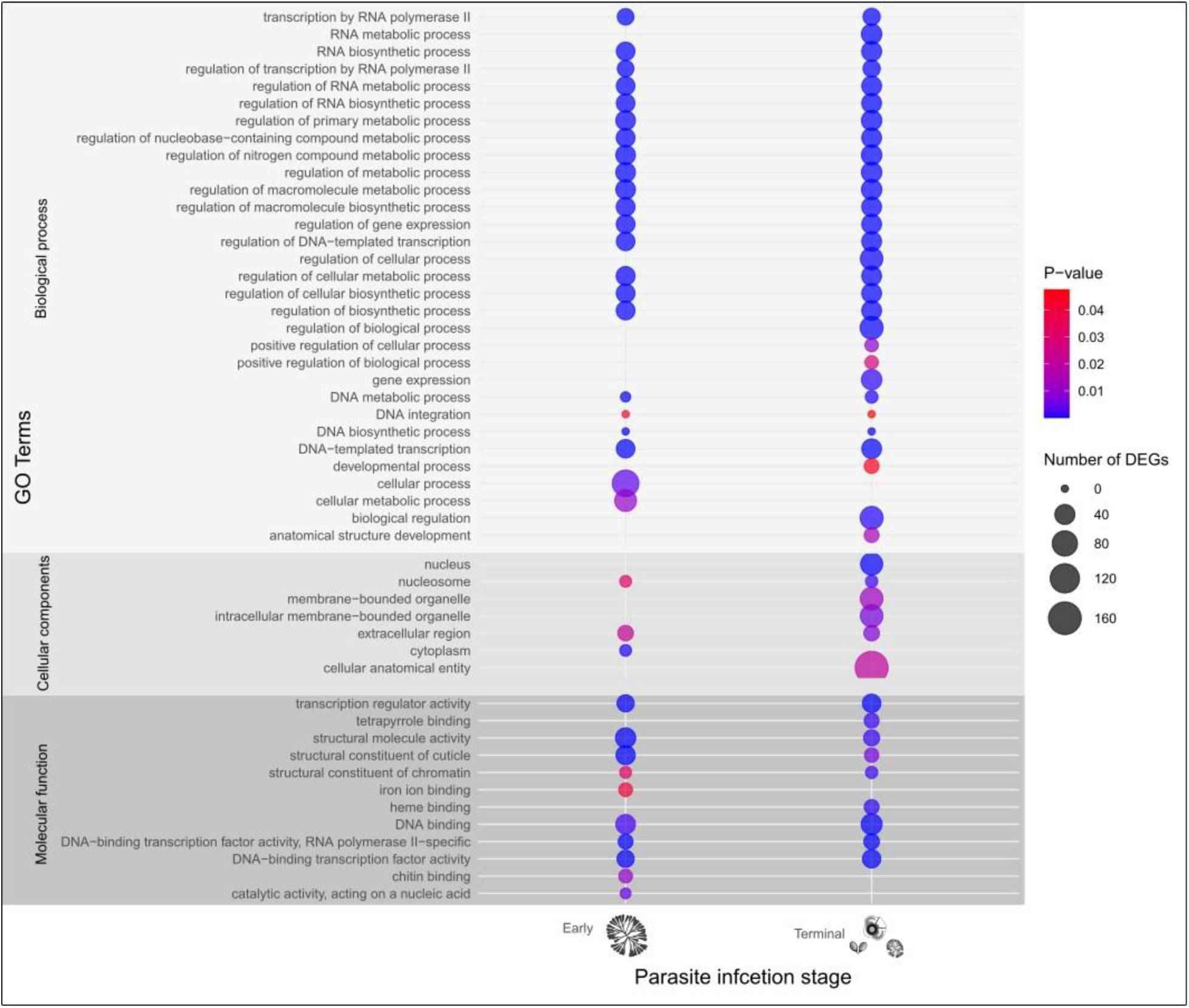
Enriched Gene Ontology (GO) terms in the Early and Terminal infection phases. The size of the circles represents the number of detected differentially expressed genes (DEGs) belonging to the respective GO term, while the color indicates the level of significance.

### Across infection phases differential pathway expression analysis

The highest number of altered pathway expression (95 pathways) was observed during the Terminal infection phase (78 upregulated, 17 downregulated), followed by Early infection (53 pathways, 36 upregulated, 17 downregulated) and Mid infection (47 pathways, 33 upregulated, 15 downregulated; Figure 4a, Tables S8-S10).

**Figure 4.**
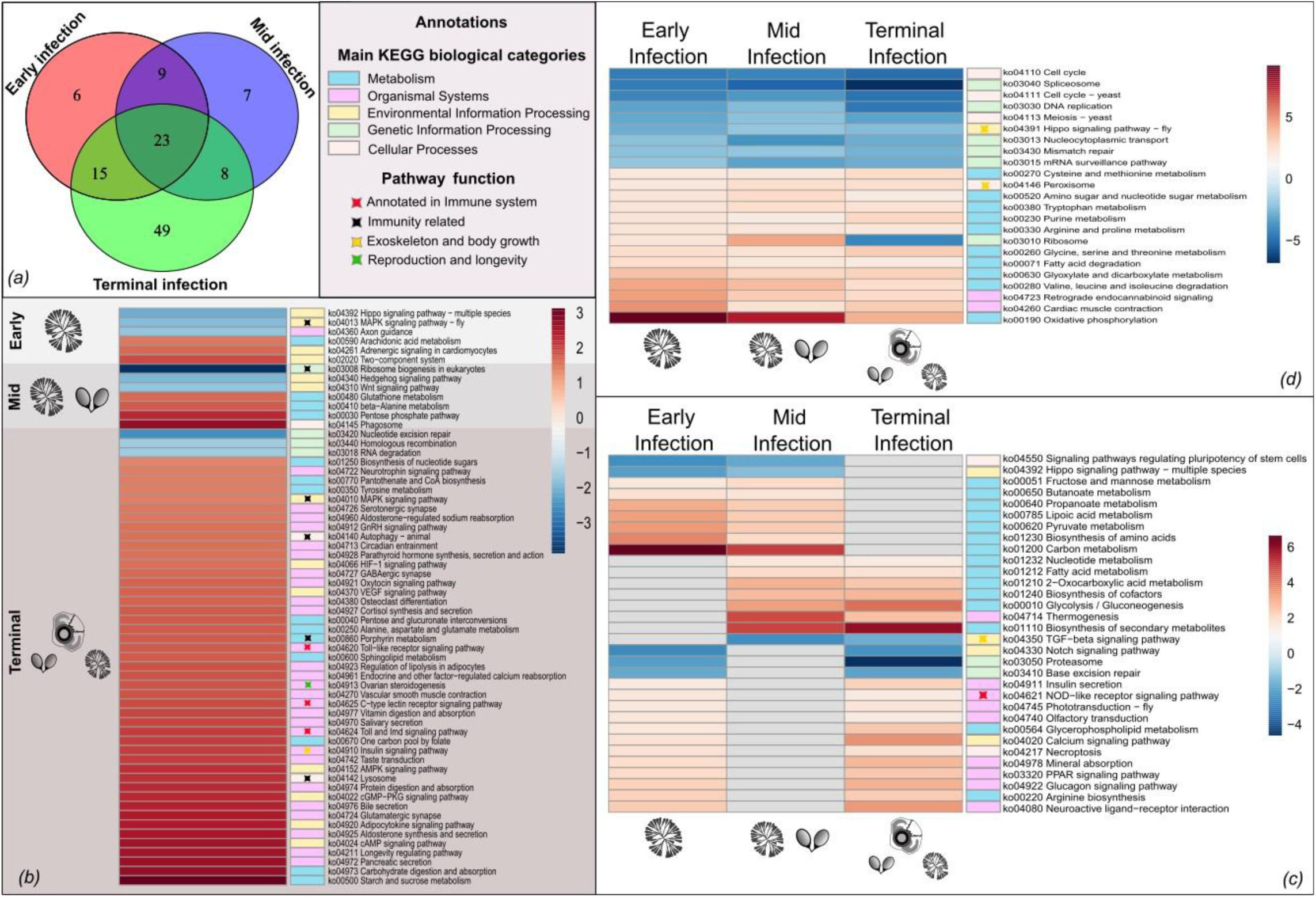
*(a)* Venn Diagram illustrating the distribution of differentially expressed pathways across infection phases. *(b)* Heatmap displaying pathways the uniquely up- and down-regulated pathways at each infection phase. *(c)* Heatmap displaying pathways that are differentially expressed in two infection phases. *(d)* Heatmap displaying pathways consistently differentially expressed across all infection phases. In all heatmaps color bars represent z-scores. Red spectrum represents upregulation in the infected individuals when compared to the controls at each respective infection phase while blue spectrum represents downregulation. Colored blocks next to the pathways represent their annotations to higher KEGG biological categories while colored asterisks indicate their likely function in respect to our findings.

During the Early infection phase, six pathways were uniquely expressed. Among them, the MAPK signaling pathway – fly (ko04013), involved in immune signaling, was downregulated (Figure 4b, Table S11). During the Mid infection phase, seven pathways were altered. Among them, the Ribosome biogenesis in eukaryotes (ko03008) pathway was downregulated (Figure 4b, Table S11), while the Phagosome (ko04145) was upregulated. Finally, during the Terminal infection phase, 46 pathways were upregulated, while three were downregulated (Figure 4b, Table S11). All downregulated pathways were related to Genetic Information Processing (Figure 4b), indicating the compromised ability of the host to repair genetic material and regulate RNA stability. Among the uniquely upregulated pathways, three pathways were annotated to immune system KEGG biological category, i.e., Toll-like receptor signaling (ko04620), C-type lectin receptor signaling (ko04625), and Toll and Imd signaling (ko04624) pathways. Additionally, pathways related to cellular damage and stress response, i.e., Autophagy (ko04140) and Lysosome (ko04142) as well as immune signaling MAPK signaling pathway (ko04010), and pathways potentially related to somatic growth involved in insulin signaling (ko04910), were upregulated in infected hosts (Figure 4b, Table S11). Pathways that might be associated with the prolonged lifespan or castration of *Daphnia* hosts during the Terminal infection phase (i.e., the Ovarian steroidogenesis associated with aging), were upregulated in infected hosts (Figure 4b, Tables S11).

There were pathways shared across two infection phases, with the majority of them annotated to either Metabolism or Organismal Systems (Figure 4c, Tables S12). The most relevant one was the NOD-like receptor signaling pathway (ko04621), which belongs to the immune system response. Additionally, pathways related to Genetic Information processing i.e., Proteasome (ko03050) and Base repair (ko03410), were downregulated. The TGF-beta signaling pathway (ko04350), likely involved in somatic growth, was downregulated in infected animals during Mid- and Terminal-infection.

Twenty-three pathways were consistently significantly-altered between infected and uninfected hosts across all the three infection phases (Figure 4d, Tables S13), likely indicating upregulated functions related to infection regardless of the infection phase. As a general pattern, pathways related to metabolism (i.e., porphyrin metabolism) and organismal systems were upregulated, while pathways related to environmental signaling, cellular processes and genetic information processing (i.e., Spliceosome) were downregulated in infected hosts. An exception to this pattern was the Peroxisome (ko04146) pathway, which was consistently upregulated. Interestingly, the Ribosome (ko03010) pathway was upregulated in Early- and Mid-infected hosts, but downregulated in Terminally infected hosts. In contrast, the oxidative phosphorylation (ko00190) pathway was highly upregulated in the Early infection phase. Additionally, the Hippo signaling pathway (ko04931), which was related before with somatic growth and development, was downregulated across all infection phases.

## Discussion

### Insights into the immune system response across infection phases

Our pathway-level expression analysis revealed enrichment of key pathways directly involved in the immune system response (Figure 4, marked with red asterisk). Specifically, the NOD-like receptor signaling pathway was upregulated in infected hosts during the Early and Terminal infection phases. NOD-like receptor signaling pathway is involved in detecting microbial components and triggering an innate immune response, thereby playing a crucial role in pathogen recognition in invertebrates (Almeida-da-Silva et al., 2023; Abdulkareem & Chen, 2021). However, in teleost fish, overactivation of NOD receptors can lead to excessive inflammatory responses, which may result in tissue damage and impaired immune function (Su et al., 2021). Therefore, the early activation of this pathway likely reflects an initial immune defense against *P. ramosa*, while its continued upregulation suggests a prolonged inflammatory response.

The Toll and Imd pathway, Toll-like receptor signaling pathway, and C-type lectin receptor signaling pathway were upregulated only during the Terminal infection phase. Regarding the Toll signaling pathways, research in *Drosophila* has demonstrated that these pathways are primarily activated in response to Gram-positive bacteria and fungi (De Gregorio et al., 2002). These pathways play a pivotal role in regulating the expression of antimicrobial peptides (AMPs), which are particularly effective against fungal infections (Hoffmann & Reichhart, 2002; Lemaitre & Hoffmann, 2007). However, *Daphnia* are not known to secrete AMPs. Given that *P. ramosa* is a Gram-positive bacterium, the upregulation of the Toll signaling response is indicative of a general immune response mechanism, rather than an effective antimicrobial defense strategy. However, the activation of those pathways during the Terminal infection phase, where the host is already colonized with mature spores, is kind of a paradox. New insights in these pathways suggest that bacterial and viral proteases subvert Toll-like receptor (TLR) signaling pathways during infectious diseases, thus evading or manipulating TLR signaling to enhance their survival and promote infection (Ciaston et al., 2022). Whether *P. ramosa* uses these specific proteases to subvert the immune system of the host is unknown. Further investigation of the *P. ramosa* genome for gene homologues of known bacterial proteases that target Toll-like receptor (TLR) signaling (e.g., Gingipains, TssM, YopJ/P, CPAF) (reviewed in Ciaston et al., 2022) could provide insights into whether this parasite actively modulates host immune signaling to evade detection and establish infection.

Interestingly, in our study, MAPK signaling, which is crucial for pro-inflammatory cytokine production, was significantly suppressed in infected hosts, a pattern also observed in bacterial infections, where proteases such as *YopJ/P* (*Yersinia*), *Lethal Factor* (*Bacillus anthracis*), and *NleD* (*Escherichia coli*) directly interfere with MAPK signaling to impair immune responses (Ciaston et al., 2022; Sweet et al., 2007). If P. ramosa uses a similar strategy, it could actively suppress host immunity by disrupting key inflammatory pathways through protease activity, thereby facilitating its establishment and persistence of infection.

### Insights into the general immune response across infection phases

Our gene- and pathway-level expression analyses revealed upregulation of key genes and pathways likely relevant to the immune response, that were not annotated to immune system (Figures 3 and 4, marked with black asterisk). Specifically, during the Terminal infection phase, infected hosts upregulated genes that encode for Cytochrome P450 proteins. Cytochrome P450 enzymes, a superfamily of heme-containing monooxygenases genes, primarily function in detoxification and metabolism (Estabrook, 2003; McDonnell & Dang, 2013), suggesting that their upregulation might be related to an inflammatory response. In parallel, genes encoding Hemoglobin (heme binding, GO:0020037) and those involved in heme biosynthesis were also upregulated. Hemoglobin is a heme-containing protein, and heme is an iron-containing porphyrin which is essential for Hemoglobin’s function, as it binds to oxygen for transport. When oxygenated, Hemoglobin becomes red due to the binding of oxygen to the iron in the heme group. The observed upregulation of genes encoding hemoglobin in infected hosts is expected to result in increased hemoglobin synthesis, which is known to contribute to the red-brownish coloration observed during this infection phase (Ebert et al., 2016).

Additionally, hemoglobin might be involved in the immune response. By channeling heme into Hemoglobin, the host could be sequestering iron and limiting the availability of free heme, a strategy that it is known as nutritional immunity (Buchmann, 2014; Iatsenko et al., 2020; Núñez et al., 2018). Pathogens such as *P. ramosa* rely on their hosts for iron, a nutrient they cannot synthesize themselves (Choby & Skaar, 2016). Therefore, they have evolved mechanisms to acquire heme from host sources, particularly Hemoglobin. For example, many bacteria employ an iron-regulated surface determinant (lsd) system where proteins on the bacterial surface bind to Hemoglobin and NEAT-containing surface proteins shuttle heme to lsd proteins to transfer them across the cell membranes. Imported heme can be incorporated into bacterial proteins, accumulated in the membrane, or degraded by heme oxygenase to release iron (Choby & Skaar, 2016). Imported lsd-mediated heme acquisition is vital for the virulence of bacteria like *Staphylococcus aureus* (Pishchany et al., 2014). However, whether *P. ramosa* has developed such exploitation systems is yet unknown. Further exploration of bacterial pathway upregulation will facilitate a better understanding of this mechanism in the *Daphnia-Pasteuria* system and enhance the comprehension of host-parasite coevolution.

Infected hosts undergo dynamic metabolic and cellular changes, balancing immune defense with parasite-induced manipulation. Host defenses rely on key degradation and recycling pathways, including Peroxisome, Lysosome and Autophagy. While Lysosome and Autophagy were specifically upregulated in the Terminal infection phase, Peroxisome activity remained consistently elevated, suggesting a sustained role in cellular homeostasis and metabolic adaptation. Autophagy acts as a cellular cleaner by encapsulating and transporting malfunctioning proteins and organelles to lysosomes for degradation (Yin et al., 2016). This mechanism plays a crucial role in regulating cell remodeling, following damage or death, in both vertebrates and invertebrates (Song et al., 2020; Moreno-Blas et al., 2025). Its activation at the Terminal infection phase may reflect that defense against pathogens remains active, even in individuals where the parasite is present in its later phases of development.

At the same time, we detected a progressive collapse in key cellular functions, which suggests likely host exhaustion. During the Mid infection phase, ribosome biogenesis was severely downregulated, yet existing ribosomes remained functional, suggesting a host stress response aimed at sustaining protein synthesis under damage, a pattern observed in invertebrates under both biotic and abiotic stress (Oakley et al., 2017; Paraskevopoulou et al., 2020; Cheng-Guang & Gualerzi, 2021). However, this compensatory mechanism failed during the Terminal infection phase, where ribosome-related pathways, along with proteasome activity, DNA repair mechanisms (mismatch repair, DNA replication), and mRNA surveillance, became strongly downregulated. This likely suggest that host carrying mature spores have reached metabolic exhaustion.

### Insights into parasite-mediated changes in exoskeleton and body growth

The “Chitin binding, GO:0008061” GO-term consisted of genes encoding for cuticular proteins, endocuticle glycoproteins, N-acetylglucosaminyltransferases and chitin-based proteins. Chitin-binding proteins, such as various cuticular proteins and chitin, a polysaccharide composed of repeating units of N-acetylglucosamine, serve as foundational components of the exoskeleton (Liu et al., 2019; Yang & Fukamizo, 2019). The upregulation of these genes can be associated with a higher frequency of molting, a process during which *Daphnia* replaces its old exoskeleton with a newer and softer one as part of its growth. Interestingly, Duneau & Ebert (2012) found that the likelihood of successful parasite infection is greatly reduced if the host molts within 12 hours after parasite exposure. This is because the attached spores are shed with the cuticle during molting, making it a beneficial response for the host when exposed to the parasite (Duneau & Ebert, 2012). Molting can thus be considered an immune defensive mechanism, emphasizing the importance of the cuticle and its development as part of the invertebrate immune response. Our results align with observations showing a significant upregulation of cuticle development-related genes in *Daphnia* exposed to a yeast pathogen (Terrill Sondag et al., 2023). The involvement of cuticle-related genes as a defensive mechanism in invertebrates has been reported previously. For instance, cuticle proteins play a pivotal role in the immune responses of insects, where host hemocytes create a melanized capsule around a pathogen to restrict its proliferation (Fedorka et al., 2013; Nappi & Christensen, 2005). Our findings, however, contradict those of Lu et al. (2018) and McTaggart et al. (2015), where proteins related to chitin were found to be downregulated in response to *P. ramosa* exposure in *D. galatea* and *D. magna*, respectively. In these studies, no upregulation or downregulation of other cuticle-related proteins was observed. Since there are two types of chitin synthases, one responsible for the synthesis of cuticular chitin and the other associated with the gut peritrophic matrix (Arakane et al., 2005; Kumar et al., 2008), we argue that in the above studies, chitin downregulation was more related to gut peritrophic matrix than to the exoskeleton.

During the Terminal infection phase, infected *Daphnia* hosts exhibited gigantism, characterized by an increase in body size, likely due to the parasite’s manipulation of host resource allocation (Ebert et al., 2004). This results in the prioritization of somatic growth over reproduction, benefiting parasite proliferation (Ebert et al., 2016). In support of this, we observed a downregulation of the Hippo and TGF-beta signaling pathways, both of which play crucial roles in growth regulation and developmental homeostasis. The Hippo pathway functions as a key suppressor of cell proliferation, and its downregulation has been linked to excessive tissue growth in model organisms like *Drosophila melanogaster* (Halder & Johnson, 2011). Similarly, TGF-beta signaling is involved in developmental patterning and tissue maintenance, and its suppression may further contribute to unregulated somatic growth. Given that infected hosts become castrated, the energy that would typically be allocated to reproduction may instead be redirected toward body size expansion. This shift could benefit *P. ramosa* by enhancing its transmission potential, as larger hosts provide a more favorable environment for parasite development and spore production, as it is implied by the *temporal storage hypothesis* (Ebert et al., 2004). According to this hypothesis, parasites can accumulate and persist within long-lived or slow-dying hosts, effectively using them as a temporal reservoir until conditions favor transmission. Thus, the observed downregulation of Hippo and TGF-beta signaling may represent a parasite-driven disruption of host growth regulation, leading to infection-induced gigantism in *Daphnia*.

Another pathway likely involved in body size changes is the “Insulin signaling pathway”. This pathway is initiated by the secretion of insulin-like peptides (ILPs), and it is essential for controlling body size, as overexpression of ILPs in invertebrates leads to increased body size by enlarging both cell size and number (reviewed in Brogiolo et al., 2001; Nijhout, 2003; Nijhout & Grunert, 2002; Oldham et al., 2002). These insulin-like molecules function as growth factors, therefore, it is possible that upregulation of insulin-related pathways contributes to excessive body growth in infected *Daphnia* during the Terminal infection phase. Parasites often manipulate host physiology, targeting key regulatory systems, including endocrine pathways. For instance, the malaria parasite *Plasmodium* mimics insulin in mosquito vectors, affecting their feeding behavior and parasite transmission (Schofield & Hackett, 1993; Sharma et al., 2019). *Schistosoma mansoni*, a parasitic flatworm, releases long non-coding RNAs (lncRNAs) that can modulate host immune responses and metabolic pathways, potentially impacting insulin signaling (Silveira et al., 2023). *Daphnia* possesses four insulin-like growth factor (IGF) receptors, suggesting sensitivity to insulin-related signals (Boucher et al., 2010). Hence, *P. ramosa* could potentially hijack the insulin secretion system in *Daphnia*, leading to increased insulin production and gigantism. However, this mechanism in bacteria has not yet been demonstrated and whether *P. ramosa* secretes insulin-like peptides or lncRNAs remains unclear. Further research is needed to reveal the molecular intricacies of the *Daphnia*-*Pasteuria* model host-parasite system.

### Insights into parasite-mediated castration

One consequence of *Pasteuria* infection in *Daphnia* is host castration, strategically diverting resources from reproduction to enhance survival, leading to the cessation of reproduction (Ebert et al., 2016). The KEGG-pathway analysis revealed an upregulation of ovarian steroidogenesis during the Terminal infection phase in infected animals. Ovarian steroidogenesis, a process whereby ovarian cells produce essential hormones to maintain reproductive tissue and trigger ovulation (Jamnongjit & Hammes, 2006), was heightened despite the castration effect. The upregulation of ovarian steroidogenesis during the Terminal infection phase, when the host is already castrated, could possibly explain why some *Daphnia* produce a final clutch towards the end of their lives, despite being castrated (i.e., castration relief; Clerc et al., 2015; Ebert et al., 2016).

## Conclusions

This study has significantly advanced our understanding of *Daphnia* host response upon infection with the bacterium *P. ramosa*. Our study highlights potential molecular mechanisms that underlie key phenotypic traits commonly observed in these infections, such as gigantism and the development of red coloration in the body cavity of the host. The analysis of gene expression across different infection phases has revealed critical functions related to immune regulation, with the most notable being the heme binding mechanism, employed as a defense strategy to limit the availability of iron, and consequently oxygen, to the parasite. It is worth mentioning that a substantial portion of the significantly enriched GO terms does not have a direct link to immunity, and many differentially expressed genes lack functional annotations. Therefore, further research should focus on annotating these uncharacterized genes to deepen our understanding of the immunological response in invertebrates.

## Author Contributions

SG, SP and JM collected and analyzed the data. SG and SP interpreted the data and wrote the manuscript with input from FBA. SG, SP, and FBA conceptualized the study. FBA obtained funding for this study.

## Acknowledgements

We are grateful to Dieter Ebert and Mariko Dale for sharing with us information that helped us formulate our experimental design. We also thank Hila Kobo, Director of the Rosalie and Harold Rae Brown Cancer Research Core Facility at the George S. Wise Faculty of Life Sciences, Tel Aviv University, for helping with the Qubit and Tape Station

## Data Availability Statement

All transcriptomic data are submitted to the Genbank Sort Read Archive (SRA) with a BioProject number (PRJNA1243082) and will be available upon publication.

